# Conserved DNA polymorphisms distinguish species in the eastern North American white oak syngameon: Insights from an 80-SNP oak DNA genotyping toolkit

**DOI:** 10.1101/602573

**Authors:** Andrew L. Hipp, Alan T. Whittemore, Mira Garner, Marlene Hahn, Elisabeth Fitzek, Erwan Guichoux, Jeannine Cavender-Bares, Paul F. Gugger, Paul S. Manos, Ian S. Pearse, Charles H. Cannon

**Affiliations:** The Morton Arboretum, Herbarium, 4100 Illinois Route 53, Lisle, IL 60532, USA; The Field Museum, Department of Botany, Chicago, IL 60605, USA; U.S. National Arboretum, Washington, DC 20002, USA; Universität Bielefeld, Department of Computational Biology, 33615 Bielefeld, GERMANY; BIOGECO, INRA, Univ. Bordeaux, 33610 Cestas, France; University of Minnesota, Minneapolis MN 55455, USA; Maryland Center for Environmental Science, Appalachian Laboratory, Frostburg, MD 21532, USA; Duke University, Durham NC 27708, USA; U.S. Geological Survey, Fort Collins Science Center, Fort Collins, CO 80526, USA

**Keywords:** Cohesion species, DNA genotyping toolkit, hybridization, introgression, *Quercus alba*, *Quercus bicolor*, *Quercus macrocarpa*, *Quercus stellata*, single nucleotide polymorphism (SNP), syngameon

## Abstract

The eastern North American white oaks, a complex of approximately 16 potentially interbreeding species, have become a classic model for studying the genetic nature of species in a syngameon. Genetic work over the past two decades has demonstrated the reality of oak species, but gene flow between sympatric oaks raises the question of whether there are conserved regions of the genome that define oak species. Does gene flow homogenize the entire genome? Do the regions of the genome that distinguish a species in one part of its range differ from the regions that distinguish it in other parts of its range, where it grows in sympatry with different species? Or are there regions of the genome that are relatively conserved across species ranges? In this study, we revisit seven species of the eastern North American white oak syngameon using a set of 80 SNPs selected in a previous study because they show differences among, and consistency within, the species. We test the hypothesis that there exist segments of the genome that do not become homogenized by repeated introgression, but retain distinct alleles characteristic of each species. We undertake a rangewide sampling to investigate whether SNPs that appeared to be fixed based on a relatively small sample in our previous work are fixed or nearly fixed across the range of the species. Each of the seven species remains genetically distinct across its range, given our diagnostic set of markers, with relatively few individuals exhibiting admixture of multiple species. SNPs map back to all 12 *Quercus* linkage groups (chromosomes) and are separated from each other by an average of 7.47 million base pairs (± 8.74 million bp, s.d.), but are significantly clustered relative to a random null distribution, suggesting that our SNP toolkit reflects genome-wide patterns of divergence while potentially being concentrated in regions of the genome that reflect a higher-than-average history of among-species divergence. This application of a DNA toolkit designed for the simple problem of identifying species in the field has an important implication: the eastern North American white oak syngameon is composed of entities that most taxonomists would consider “good species,” and species in the syngameon retain their genetic cohesion because characteristic portions of the genome do not become homogenized despite a history of introgression.

---

Hybridization and introgressive gene flow in oaks have long suggested the question of what constitutes an oak species. The 1867 edition of Gray’s *Manual of the Botany of the Northern United States* (Gray, 1867), for example, reports five hybrids in oaks,^1^ and Wiegand (1935) notes that in this edition, “we find hybrids scarcely mentioned except in one genus, *Quercus*.” In the early 20^th^ century, studies of character segregation in first and second-generation oak hybrids suggested that adaptive gene flow might contribute to range extensions in the southern live oak *Quercus virginiana* (Ness, 1918; Allard, 1932; Yarnell & Palmer, 1933). The roughly 100 years following Gray’s 1867 edition saw a number of seminal papers, mostly dealing with the effects of interspecific hybridization on oak species origins, coherence and evolutionary trajectories (e.g., Engelmann, 1876; Palmer, 1948; Muller, 1952).

In the mid 1970s, a trio of now-classic papers focused on the eastern North American white oak syngameon set the stage for contemporary studies of oak species coherence. In 1975, James Hardin published an article in the *Journal of the Arnold Arboretum* reporting evidence of widespread gene flow among 16 white oaks of eastern North America (Hardin, 1975). At about the same time, a pair of articles in *Taxon* argued that gene flow in oaks is dominated by localized gene flow among individuals that are closely enough related to exchange genes, irrespective of species, rather than among populations within species (Burger, 1975; Van Valen, 1976). Because of ongoing gene flow and introgression, Burger and Van Valen argued, oak species cannot be defined by reproductive isolation. Rather, oak species represent ecologically discrete lineages with distinct evolutionary trajectories. “Species,” Van Valen wrote, “are maintained for the most part ecologically, not reproductively.” He and Burger both argued that local gene flow among sympatric populations of different species may exceed gene flow between geographically distant populations of single species, and that the capacity for interbreeding cannot therefore be the criterion by which we recognize oak species. Burger went so far as to suggest erecting subgenera or sections that are equivalent to reproductive species, but allowing our named species in oaks to represent ecologically and morphologically defined evolutionary lineages. The idea that gene flow is often insufficient to cause species to cohere across their range had been discussed previously (Ehrlich & Raven, 1969), but Burger and Van Valen seem to be making a stronger claim: oak species are delimited not reproductively, but ecologically. A measured skepticism about oak species is not uncommon among botanists even today, unsurprising in the face of ample evidence of introgression and gene flow (e.g., Whittemore & Schaal, 1991; Dumolin-Lapegue *et al.*, 1997; Dumolin-Lapegue, A., & Petit, 1999; Petit *et al.*, 2003; Dodd & Afzal-Rafii, 2004; Tovar-Sánchez & Oyama, 2004; Craft & Ashley, 2006; Lexer, Kremer, & Petit, 2006; Curtu, Gailing, & Finkeldey, 2007; Hipp & Weber, 2008; Chybicki & Burczyk, 2010; Moran, Willis, & Clark, 2012).

In the past two decades, the increased availability of single-locus DNA markers has stimulated investigation into the processes that maintain distinct species in the presence of interspecific hybridization (Kremer & Hipp, Accepted pending revision). It is notable that different studies using single-locus DNA markers have shown strikingly different patterns. Studies utilizing chloroplast DNA markers have generally yielded clear evidence of introgressive exchange of markers, with little if any clustering of individuals by species (Whittemore & Schaal, 1991; Dumolin-Lapegue *et al.*, 1997, 1999; Petit *et al.*, 1997, 2003; Manos, Doyle, & Nixon, 1999; Belahbib *et al.*, 2001; Pham *et al.*, 2017). Studies utilizing nuclear markers, on the other hand, have typically demonstrated that gene flow among species (Dodd & Afzal-Rafii, 2004; Gömöry & Schmidtová, 2007; de Casas *et al.*, 2007; Eaton *et al.*, 2015) is balanced by gene flow within species, promoting species cohesion (Whittemore & Schaal, 1991; Muir, Fleming, & Schlötterer, 2000; Muir & Schlötterer, 2005; Lexer *et al.*, 2006; Leroy *et al.*, 2017, 2018).

Next generation DNA sequencing (NGS) has made it practical to test more rigorous models of introgression history in oaks using much larger numbers of loci (e.g., Eaton *et al.*, 2015; Leroy *et al.*, 2017). Additionally, NGS has enabled economical development of genotyping toolkits for smaller applications. In a recent paper, we utilized a large RAD-seq dataset for white oaks (McVay, Hipp, & Manos, 2017b; Hipp *et al.*, 2018) to develop a low-cost SNP genotyping kit for eastern North American white oaks (Fitzek *et al.*, 2018). We demonstrated our 80-marker SNP kit to be effective for identifying 15 species and F_1_ hybrids, and validated it in a garden setting, where we found hybridization between non-native species in the collection and the native white oaks of the surrounding woodlands. In the current study, we test this marker set in natural populations across a rangewide sample of seven eastern North American white oaks. These species are components of a classic syngameon, where there is good documentation of interspecific hybridization in many combinations (Hardin, 1975) and introgressive exchange of chloroplast haplotypes (Whittemore & Schaal, 1991; Pham *et al.*, 2017). We investigate whether the species are genetically cohesive at these 80 loci or a subset thereof, representing areas of the genome that have presumably been shielded from introgression across the range of the species. We also map these markers back to a chromosome-level assembly of the *Quercus robur* L. genome (Plomion *et al.*, 2018) to investigate whether they are distributed across the genome or, conversely, whether genetic cohesion of the eastern North American white oaks is concentrated in a few genomic islands of differentiation. Our study provides a first framework investigation of the eastern North American white oak syngameon using a genomewide sample of molecular markers, laying the groundwork for future studies of introgression and species cohesion in the group.

## Materials and methods

### Sampling and genotyping

Data were initially collected from 184 individuals of seven eastern North American white oak species, collected from a wide geographic range for each species; in this study, *Quercus muehlenbergii* Engelm. and *Q. prinoides* Willd. are separated in name only, as our RAD-seq data failed to distinguish the species (McVay *et al.*, 2017b; Hipp *et al.*, 2018) and SNPs were consequently not designed to separate these two (Fitzek *et al.*, 2018). The species status of these two bears investigation with broader sampling. Throughout the remainder of this paper, we will refer to these two together as *Q. muehlenbergii / prinoides*, not because we are making a claim that they are not distinct taxonomically, but to reflect the fact that they are grouped for analysis. Samples represent unique adults with seven exceptions, for which a second extraction of each individual was genotyped as a technical replicate. Individuals were selected to be typical of the species morphologically, not to be a random sample of all potential pure and introgressed individuals. Twenty-one individuals for which fewer than 90% of loci amplified successfully were removed from analysis and are not discussed further in this paper, leaving a final set of 163 individuals analyzed (Fig. 1; Table 1).

**Figure 1.**
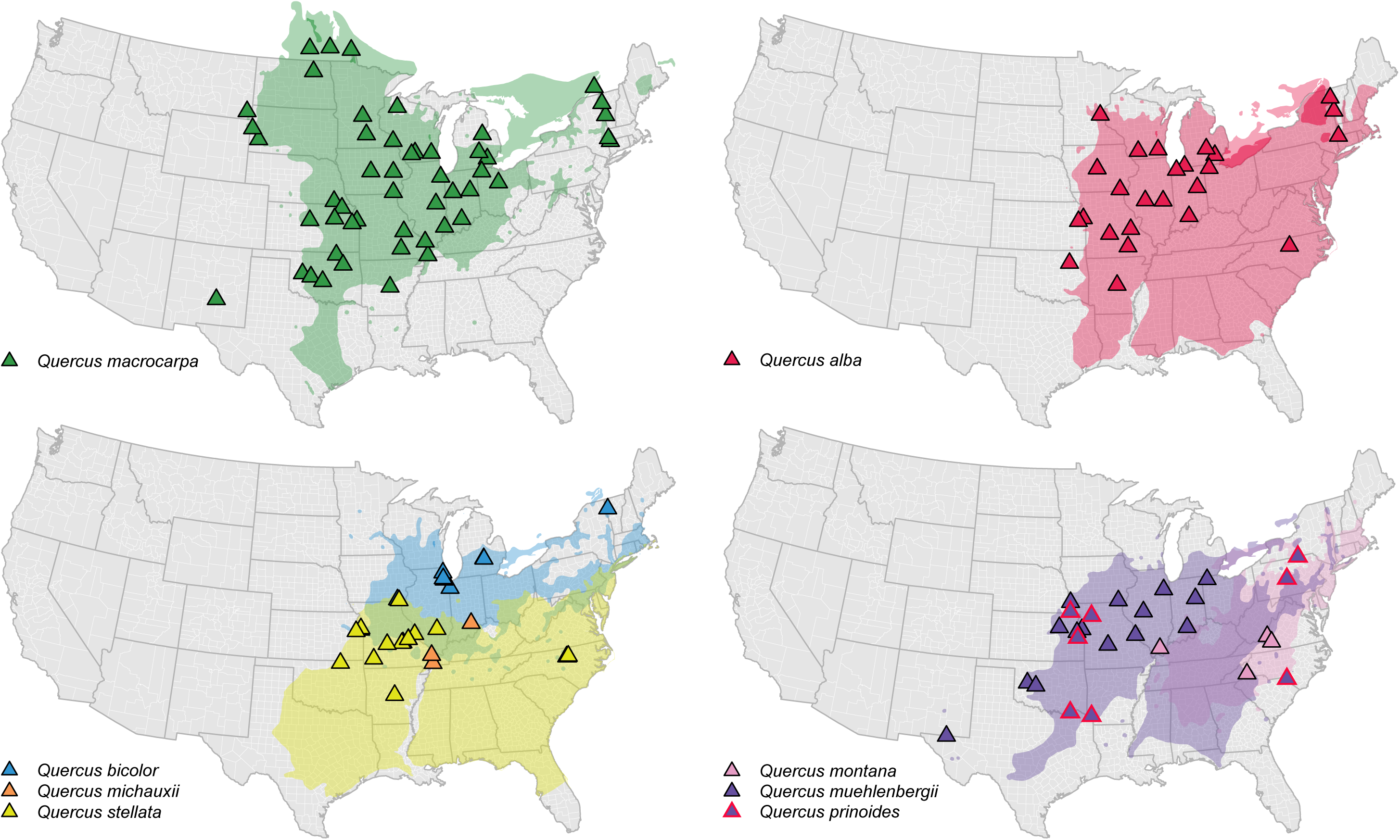
Sampling map. Sites were sampled to roughly cover the range of the taxa as known; on each panel, collections are overlaid on the range maps for each species following Little (1971, 1977, 1979), except for *Quercus prinoides*, for which a base map was not available.

**Table 1.**
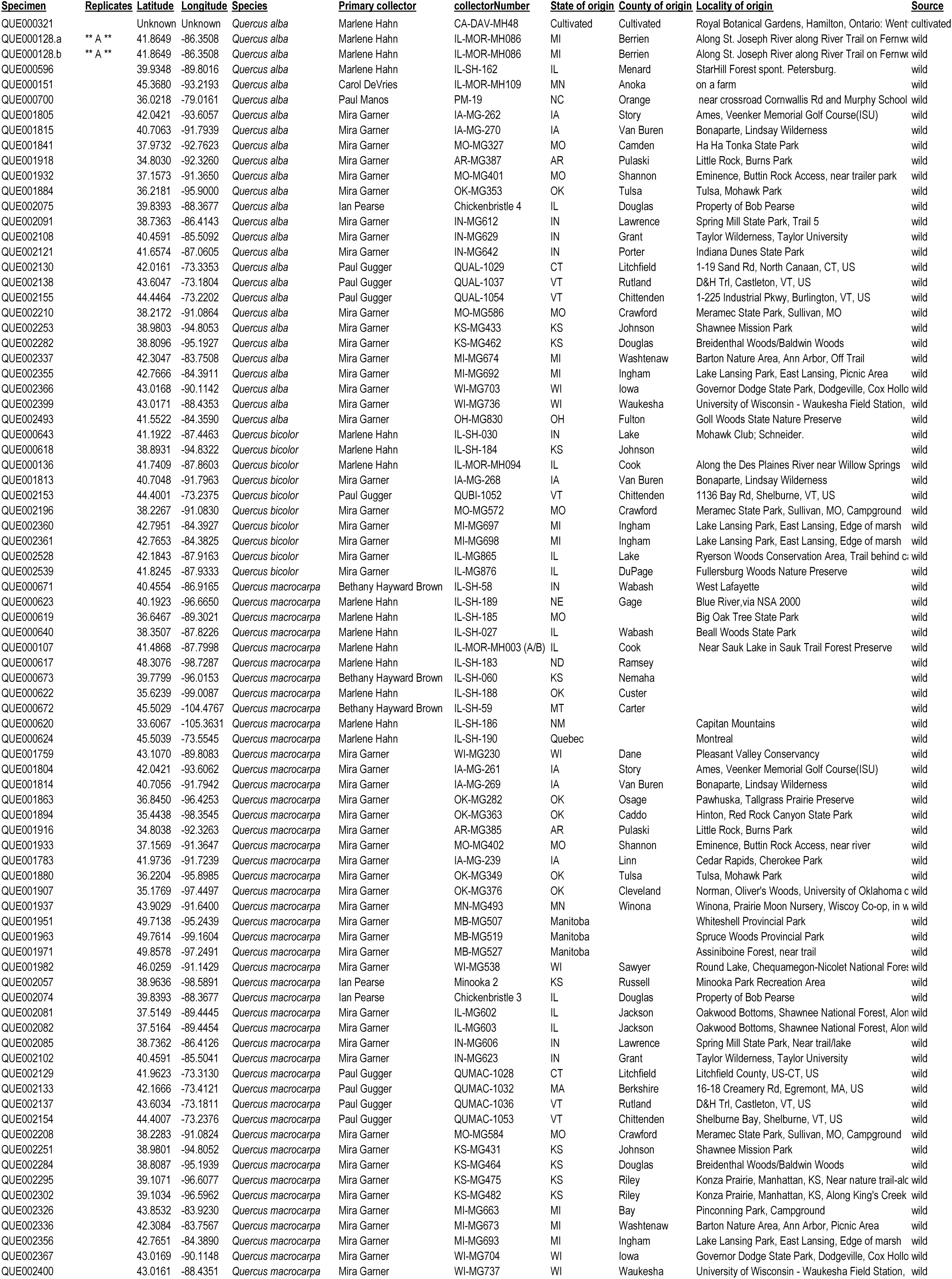
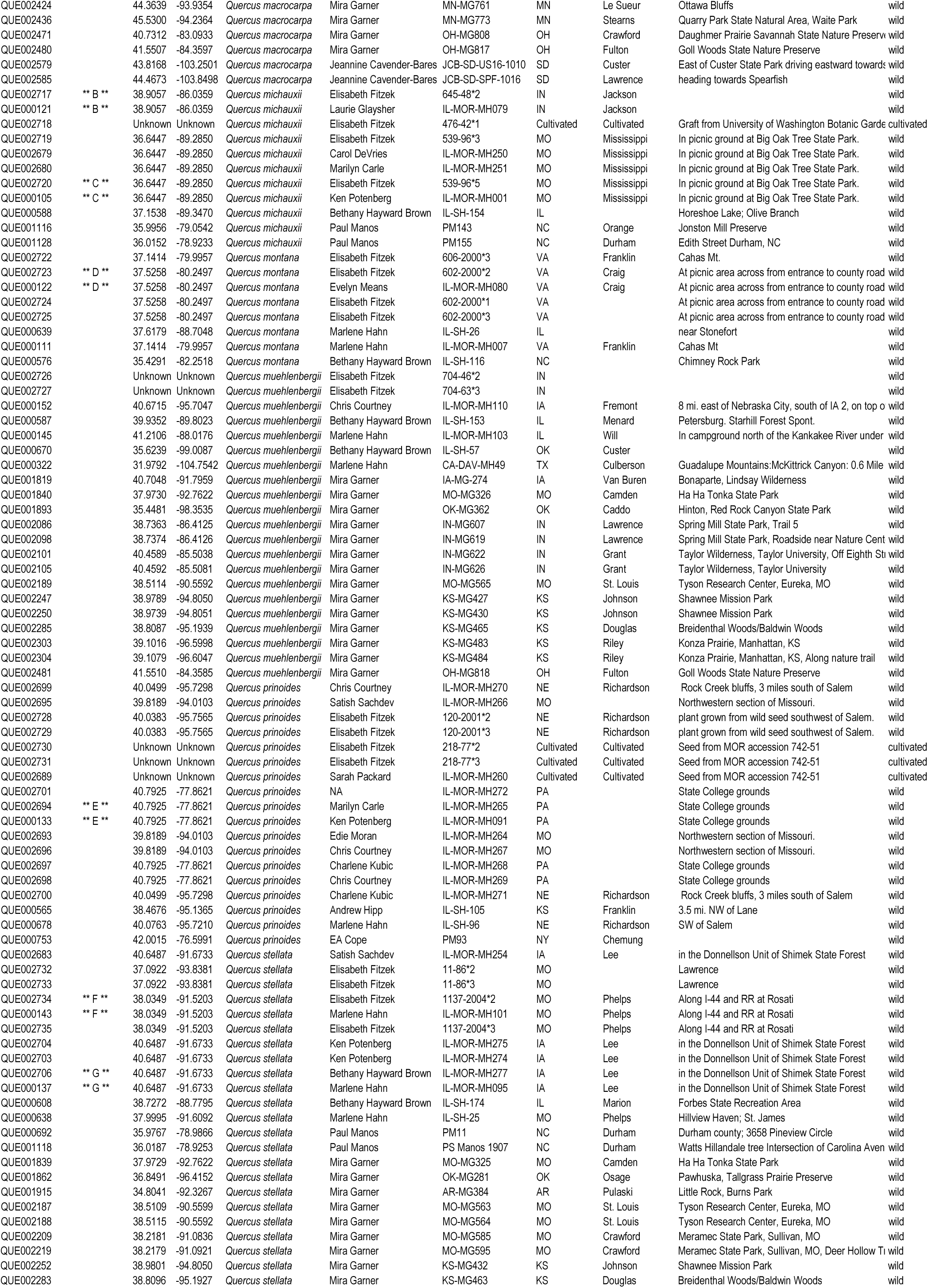
Samples included in study. Locality and coordinate data indicate source populations for both wild and cultivated material; where material is of cultivated source, no state or county data are provided. Replicates indicate technical replicates extracted from the same individual: individuals with the same replicate code are identical. *[note to editors:* Table 1 *was provided as PDF and XLSX; please format for inclusion in text]*

To reduce the opportunity for hybridization with taxa from outside the natural range of each species, samples were preferentially selected from wild populations or from trees grown in gardens from seeds of known wild provenance (as discussed in Fitzek *et al.*, 2018; Hipp *et al.*, 2018); five individuals were analyzed from cultivated material (Table 1). Sample size per species ranges from 7–9 in *Quercus montana* Willd. and *Q. michauxii* Nutt. to 38–52 in *Q. muehlenbergii / prinoides* and *Q. macrocarpa* respectively (Table 1). The distance between the most widely separated populations sampled within each species ranges from 771 km in *Q. montana* to 3005 km in *Q. macrocarpa* (Table 2). Moreover, aside from samples of *Quercus macrocarpa* at the westernmost and northernmost edges of its range (Fig. 1), almost all samples in our study were collected from within the range of at least one other species. Consequently, while our study does not encompass the entire range of each species, the samples cover a wide geographic range within each species, with the opportunity for crossing among congeners. Locations for source populations of all samples for which source information was available were plotted over range maps for *Q. macrocarpa*, the most wide-ranging species in our study; *Q. bicolor*, the most widespread northern species; and *Q. stellata*, the most widespread southern species. Range maps were plotted from shapefiles (Prasad & Iverson, 2003) generated from previously published range maps of North American trees (Little, 1971, 1977, 1979) over the ‘county’ and ‘state’ base maps provided in maps v. 3.3.0 (Becker *et al.*, 2018) for R v. 3.4.2, ‘Short Summer’ (R-Development-Core-Team, 2004). All plotting was done in R using the ggplot2 (Wickham, 2009) and ggmap (Kahle & Wickham, 2013) packages, using proj4 (Urbanek, 2012) for map projections.

**Table 2.** Sample sizes, sample distances and ranges, and overall species ranges. Sample distance (D) maximum and median were calculated from Table 1 using the Haversine formula. Species ranges were inferred from range maps of Little (1971, 1977, 1979) for all species except *Q. prinoides*, which was estimated by visual inspection of maps published in Flora of North America (Nixon, 1977). *[note to editors:* Table 2 *was provided as PDF and XLSX; please format for inclusion in text]*

|  | <u>N</u> | <u>Sample D max (km)</u> | <u>Sample D median (km)</u> | <u>Sample latitude</u> | <u>Species latitude</u> | <u>Sample longitude</u> | <u>Species longitude</u> |
| --- | --- | --- | --- | --- | --- | --- | --- |
| *Quercus macrocarpa* | 52 | 3005.3 | 888.8 | 33.6, 49.9 | 28, 52.7 | -105.4, -73.2 | -104.4, -66.1 |
| *Quercus alba* | 26 | 2120.1 | 695 | 34.8, 45.4 | 29.6, 46.5 | -95.9, -73.2 | -96.3, -69.1 |
| *Quercus muehlenbergii* | 21 | 2098.3 | 543 | 32, 41.6 | 24.8, 44.7 | -104.8, -84.4 | -105.2, -72.2 |
| *Quercus stellata* | 21 | 1565.6 | 325.6 | 34.8, 40.6 | 27.6, 41.8 | -96.4, -78.9 | -101.4, -70 |
| *Quercus prinoides* | 17 | 1618.9 | 185.9 | 38.5, 42 | 34.1, 42.9 | -95.8, -76.6 | -99.8, -70 |
| *Quercus bicolor* | 10 | 1889.8 | 453.4 | 38.2, 44.4 | 35.2, 46.4 | -94.8, -73.2 | -96.4, -70 |
| *Quercus michauxii* | 9 | 939 | 380.5 | 36, 38.9 | 28.8, 41 | -89.3, -78.9 | -95.5, -74.3 |
| *Quercus montana* | 7 | 771.3 | 277.8 | 35.4, 37.6 | 32, 44.6 | -88.7, -80 | -90, -70.5 |

Samples were genotyped using an 80-SNP DNA toolkit developed to distinguish 15 eastern North American white oaks (as described in Fitzek *et al.*, 2018). Briefly, an extensive RAD-seq dataset comprising multiple exemplars of all 15 species (McVay *et al.*, 2017b) was surveyed for SNP variation, using pairwise F_ST_ to identify SNPs that were (1) fixed or nearly fixed between species and (2) flanked by at least 20 bp of conserved sequence, which could be used for primer design. Multiplexes of up to 40 primers for potential SNPs were designed using the Assay Design 4.0 Suite (Agena Biosciences, San Diego), which is optimized for MassARRAY analysis (Bradić, Costa, & Chelo, 2012). Samples were genotyped using the iPLEX Gold chemistry following Gabriel et al (2009) on a MassARRAY system (Agena Biosciences) at the Genomic Platform of Bordeaux with the help of Adline Delcamp. Data analysis was completed using MassARRAY Typer Analyzer 4.0.26.75 (Agena Biosciences). We manually checked each marker clustering to detect potential ambiguous genotype assignation or unusable SNP. The results were exported as a genotype table for downstream analyses. After genotyping, 5 SNPs were removed from analysis because they failed to amplify in more than 30% of individuals.

The oak genome was not yet available when this DNA toolkit was published, but since then a chromosome-level genome has been published for *Quercus robur* (Plomion *et al.*, 2018), a white oak closely related to the species for which this toolkit was developed. To evaluate the genomic independence of the loci we used in this study, all RAD-seq loci used to develop the 80 SNPs were mapped to the oak genome using BLASTN (Altschul *et al.*, 1990; Camacho *et al.*, 2009) with a threshold EValue of 0.0001. Each RAD-seq locus was identified as mapping to a single position on a chromsome, multiple positions, or not mapping. All SNPs were designed from distinct RAD-seq loci save two (CL_2457_66 and CL_2457_32), which both come from a single RAD-seq locus that maps to position 36,055,433 on *Quercus robur* chromosome 12. The two SNPs identified in this RAD-seq were designed to distinguish *Quercus stellata* from the remaining taxa and should not be considered independent of one another. They are not strongly decisive and do not figure prominently in downstream analyses in this study or in Fitzek et al. (2018).

Genomic clustering of loci was evaluated by calculating intervals between loci on each chromosomes and comparing these to a simulated null distribution. The null distribution was simulated based on 10,000 replicate datasets of 59 loci drawn at random from the 41,898 uniquely mapped *Pst*I RAD-seq loci from the larger study from which our SNPs were developed (Hipp *et al.*, 2019). Three test statistics were evaluated: mean interval length between all loci on all chromosomes; number of intervals < 1E04 bp; and number of intervals < 1E06 bp. Code for performing this test is archived in https://github.com/andrew-hipp/white-oak-syngameon.

### Data analysis: evaluating species cohesion

We define species cohesion operationally in this study using two criteria: (1) clustering of all plants sampled from each species in genetic space, exclusive of other species, and irrespective of geography; and (2) minimal evidence of genetic admixture between species at some conserved region of the genome (in this case, based on preselected markers). By this definition, clustering of individuals by geography instead of by species would be evidence against species cohesion, as would any proportion of the genome of individuals of a putative species that is shared with individuals of other putative species. This operational definition corresponds with practices widely used by plant systematists to define “good species” (Rieseberg, Wood, & Baack, 2006) as well as statistical methods traditionally used to infer patterns and degree of interspecific introgression (Anderson, 1949). It puts off for the time being possible empirical and philosophical issues with cohesion species as a concept (Barker, 2007; Barker & Wilson, 2010) as well as questions about the mechanisms by which species cohere (Morjan & Rieseberg, 2004).

We assess criterion 1, clustering in genetic space, using the unweighted pair group method with arithmetic mean (UPGMA) (Sokal & Michener, 1958), a clustering method that aggregates individuals based on a pairwise distance matrix, in this case a Euclidean distance matrix based on allele counts within individuals, where each allele is present as 0, 1, or 2 copies per individual. UPGMA is well suited to within-species comparisons of genetic data or other comparisons of data that are truly ultrametric, where it performs reasonably well as an estimator of genetic relatedness (Felsenstein, 2004). In our study, UPGMA has the desirable property of apportioning genetic variance to branches, so that we can assess whether the variance in our data is better assigned to among-species or within-species differences. Because our markers are designed with extreme bias toward among-species differences, we do not attempt to quantify variance components using AMOVA (Excoffier, Smouse, & Quattro, 1992) and urge that the clustering results not be interpreted as estimating these variance components. We compare UPGMA results with non-metric multidimensional scaling (NMDS) ordination on the same data matrix. We present results from the three-dimensional ordination because it suffices to discriminate the species in our study.

Criterion 2 we assess using the Bayesian population genetic clustering algorithm implemented in STRUCTURE v 2.3.4 (Pritchard, Stephens, & Donnelly, 2000). We utilized the admixture model with correlated allele frequencies and λ fixed at 1.0, allowing *K* (the number of populations) to range from 1 to 12. For each value of *K*, we ran 10 replicate MCMC runs of 1E06 generations following a 1E05 generation burn-in. We followed the method of Evanno et al. (2005) to identify the most probable value of *K* based on the maximum value of Δ*K*, but given the problematic nature of identifying *K* with hierarchical data, we report the structures recovered under multiple values of *K*. We utilized STRUCTURE HARVESTER (http://taylor0.biology.ucla.edu/structureHarvester/) (Earl & vonHoldt, 2012) to calculate the Evanno statistics and CLUMPP v 1.1.2 for 64 bit Linux (Jakobsson & Rosenberg, 2007) to average STRUCTURE run replicates for each value of *K*. We visualized results using DISTRUCT v. 1.1 (Rosenberg, 2004).

To evaluate whether the entire SNP toolkit is necessary to discriminate among the species we are studying and to identify SNPs that might be fixed within species, we calculated the absolute number and proportion of individuals within each species possessing each polymorphism observed. With the caveat that sampling is uneven across species (ranging from N = 7 in *Q. montana* to N = 52 in *Q. macrocarpa*), the resulting heatmap (Fig. 2) and the table underlying it (Supplemental Table S1) estimate the decisiveness of each SNP relative to species identification in this species group: the summed proportion of individuals by species that have a given SNP estimates that SNP’s decisiveness, where a sum of 1.0 or 2.0 (for *Q. muehlenbergii / prinoides*) indicates a locus that is alone decisive for a taxon for the samples we have genotyped. The reduced set may have practical benefit for both cost and because the combinability of primer pairs plays a crucial role in multiplexing (Fitzek *et al.*, 2018). Decisiveness was overlaid on the mapped SNPs to identify whether loci that are fixed or nearly fixed within species are genomically clustered (Table 3).

**Figure 2.**
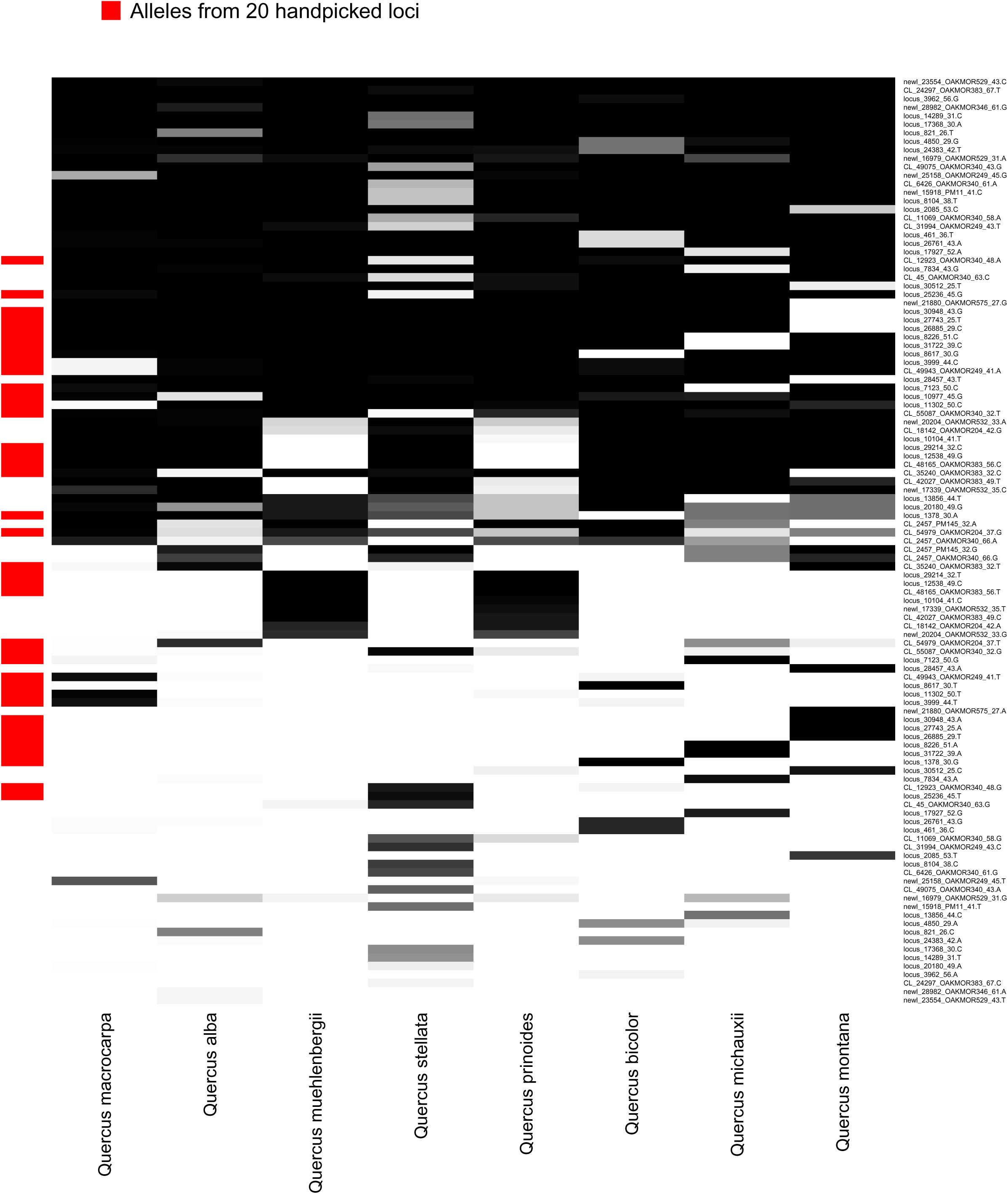
SNP heatmap by species. Darkness of cells indicates the percent of individuals of a given named species possessing the indicated nucleotide. Red bars along the side of the figure indicate SNPs in 20 loci we hand-selected because they were highly decisive for the species represented in the present study.

**Table 3.** Map positions and decisiveness of SNPs that map to a unique position on one of the *Quercus robur* chromosomes. The 60 SNPs that map back to one of the 12 *Quercus robur* chromosomes inferred in Plomion et al. (2018) are shown here with their start position on the *Q. robur* chromosome to which they map and their decisiveness, abbreviated as follows: ‘***’ indicates a SNP whose decisiveness is exactly 1.000 or 2.000 for the sample studied here (i.e., diagnostic for one or two species); ‘**’ if it is within 0.100 of 1.000 or 2.000; or ‘*’ if it is within 0.200 of 1.000 or 2.000. All loci mapped with identity > 95%, locus length > 70 bp, and E-value < 1.0 × 10^−30^. The table demonstrates that the most decisive loci in our toolkit are distributed across all chromosomes except 5, 7, and 11, and separated by an average of 7.47 million bp ± 8.74 million bp (s.d.). Four pairs of loci are < 10,000 bp from one another (indicated by bold italics in the table under “Dist. (bp)” and may bear further investigation as possible islands of differentiation. Mapping details from BLASTN and mapping information from non-uniquely mapping loci and loci that did map are in Supplemental Table S3. Abbreviations: Query (locus) = the RAD-seq locus SNP abbreviation from Fitzek et al. 2018; LG = linkage group (chromosome number), following Plomion et al. 2018; Start (bp) = start position of the RAD-seq locus on the *Q. robur* chromosome; Dist. (bp) = distance in base pairs from the start of the locus to the end of the locus adjacent to it on the same chromosome; Decisiveness = decisiveness of the SNP for identifying one species or a pair of species (cf. Fig. 2).

| <u>Query (locus)</u> | <u>LG</u> | <u>Start (bp)</u> | <u>Dist. (bp)</u> | <u>Decisiveness</u> |
| --- | --- | --- | --- | --- |
| locus_11631_48 | 01 | 1.32E+07 |  |  |
| locus_17927_52 | 01 | 1.92E+07 | 6.04E+06 | * |
| newl_21880_27 | 01 | 2.02E+07 | 9.71E+05 | *** |
| locus_821_26 | 01 | 4.61E+07 | 2.59E+07 |  |
| newl_17339_35 | 01 | 4.85E+07 | 2.40E+06 | ** |
| CL_42027_ | 02 | 9.97E+06 |  | ** |
| CL_6426_61 | 02 | 1.46E+07 | 4.62E+06 |  |
| locus_4492_52 | 02 | 2.38E+07 | 9.21E+06 |  |
| locus_20180_49 | 02 | 2.72E+07 | 3.37E+06 |  |
| locus_3962_56 | 02 | 2.72E+07 | 1.90E+03 |  |
| CL_49075_43 | 02 | 4.71E+07 | 1.99E+07 |  |
| CL_35240 | 02 | 5.65E+07 | 9.40E+06 | ** |
| locus_12538_49 | 02 | 5.65E+07 | 9.16E+03 | *** |
| locus_23517_52 | 02 | 6.66E+07 | 1.01E+07 |  |
| newl_23554_ | 02 | 7.19E+07 | 5.26E+06 |  |
| locus_3169_44 | 02 | 7.55E+07 | 3.68E+06 |  |
| locus_9121_49 | 02 | 9.24E+07 | 1.68E+07 |  |
| CL_55087_32 | 02 | 9.38E+07 | 1.43E+06 | * |
| locus_8059_35 | 03 | 2.99E+07 |  |  |
| CL_11069_58 | 03 | 2.99E+07 | 2.65E+03 | * |
| locus_8717_53 | 03 | 3.81E+07 | 8.25E+06 |  |
| locus_7123_50 | 03 | 4.03E+07 | 2.18E+06 | ** |
| locus_5882_32 | 03 | 5.25E+07 | 1.22E+07 |  |
| locus_8617_30 | 04 | 3.15E+07 |  | ** |
| locus_5229_56 | 05 | 5.16E+07 |  |  |
| locus_29214_32 | 06 | 1.28E+07 |  | *** |
| locus_10977_45 | 06 | 2.04E+07 | 7.58E+06 |  |
| CL_54979_ | 06 | 3.53E+07 | 1.49E+07 |  |
| CL_12923_ | 06 | 4.47E+07 | 9.42E+06 | ** |
| locus_7834_43 | 06 | 4.58E+07 | 1.02E+06 | ** |
| newl_27648_32 | 07 | 1.25E+07 |  |  |
| locus_5482_34 | 07 | 2.57E+07 | 1.32E+07 |  |
| locus_27412_25 | 07 | 3.67E+07 | 1.10E+07 |  |
| locus_30948_43 | 08 | 9.66E+05 |  | *** |
| locus_5422_58 | 08 | 4.43E+07 | 4.34E+07 |  |
| locus_26761_43 | 08 | 5.11E+07 | 6.78E+06 | *** |
| locus_24383_42 | 08 | 6.07E+07 | 9.62E+06 |  |
| locus_10104_41 | 08 | 6.92E+07 | 8.51E+06 | ** |
| locus_28457_43 | 09 | 1.90E+07 |  | ** |
| newl_16979_31 | 09 | 2.81E+07 | 9.05E+06 |  |
| locus_1378_30 | 09 | 3.61E+07 | 8.08E+06 | *** |
| locus_30512_25 | 10 | 2.75E+06 |  | ** |
| locus_2085_53 | 10 | 3.57E+07 | 3.29E+07 |  |
| locus_20667_37 | 11 | 2.92E+07 |  |  |
| locus_14289_31 | 11 | 3.82E+07 | 9.02E+06 |  |
| CL_48165 | 12 | 1.46E+07 |  | *** |
| locus_11302_50 | 12 | 1.61E+07 | 1.55E+06 | ** |
| locus_31722_39 | 12 | 1.75E+07 | 1.34E+06 | *** |
| locus_9837_55 | 12 | 2.05E+07 | 3.04E+06 |  |
| newl_25158_45 | 12 | 2.11E+07 | 5.67E+05 |  |
| locus_26885_29 | 12 | 2.11E+07 | 3.37E+03 | *** |
| locus_792_52 | 12 | 2.11E+07 | 5.29E+04 |  |
| locus_8226_51 | 12 | 2.11E+07 | 7.99E+03 | *** |
| locus_25236_45 | 12 | 2.12E+07 | 9.02E+04 | ** |
| newl_15918_PM11_41 | 12 | 2.41E+07 | 2.83E+06 |  |
| locus_10802_36 | 12 | 2.99E+07 | 5.81E+06 |  |
| locus_17368_30 | 12 | 3.54E+07 | 5.49E+06 |  |
| CL_2457_32 | 12 | 3.61E+07 | 6.97E+05 |  |
| CL_2457_OAK-MOR-340_66 | 12 | 3.61E+07 | *same locus* |  |
| locus_4850_29 | 12 | 3.92E+07 | 3.12E+06 |  |

All data and code required to reproduce analyses presented here are archived in https://github.com/andrew-hipp/white-oak-syngameon.

## Results

In the full dataset of 184 individuals for 80 loci, missing data per individual averaged 2.56% ± 4.10 (s.d.) loci, and missing data per locus averaged 14.6% ± 26.8 (s.d.) individuals. In the dataset cleaned to 163 individuals for 75 loci, excluding individuals with >10% missing loci and loci with > 30% missing individuals, missing data dropped to 1.19 ± 1.13% missing loci per individual and 5.60 ± 13.9% missing individuals per locus. Of the 75 cleaned loci, 20 were monomorphic and 55 had two or more polymorphisms. Among 7 pairs of technical replicates, a total of 38 differences were found. Of these, 37 were differences in whether a locus amplified or not; only one difference in allele call was found (for locus CL_55087_OAKMOR340_32, G/T in *Quercus stellata* QUE002706 vs. G/G in specimen QUE000137). Thus among 7 × 75 = 525 replicated sites, only one genotyping error (0.17%) and 37 loci that failed to amplify in one of the two replicates (6.43%) were detected.

Seven loci exhibit only a single SNP for exactly one species in our dataset—one in *Q. alba*, two in *Q. michauxii*, four in *Q. montana*—and three exhibit a single SNP in *Q. muehlenbergii / prinoides*. An additional ten SNPs exhibit a summed proportion between 0.95 and 1.05, suggesting relatively high decisiveness for *Q. stellata* (2 SNPs) and *Q. bicolor* (3 SNPs). Based on these, we hand-picked 20 SNPs that suffice to diagnose the species in our study (Fig. 2, red bars along left edge).

Using all loci, the UPGMA (Fig. 3a) and NMDS ordination (Fig. 4) both clearly separate individuals by species, except for *Quercus prinoides* and *Q. muehlenbergii*, which our SNP genotyping primers were not designed to distinguish from one another. Thus there are seven distinct clusters recognized in this study. Individuals of these clusters separate with no overlap in three dimensional genetic ordination space (Fig. 4; note that while some species overlap in one or two dimensions, none overlap in all three) and UPGMA stem lengths that equal or exceed the species crown depth for four of the clusters (*Q. macrocarpa*, *Q. bicolor*, *Q. muehlenbergii / prinoides*, and *Q. montana*) and, for the other three, stem lengths that are approximately equal to (*Q. stellata*, *Q. michauxii*) or substantially less than (*Q. alba*) the crown height. Using the 20 hand-picked loci, our SNP genotyping toolkit successfully distinguishes species from one another using UPGMA (Fig. 3b).

**Figure 3.**
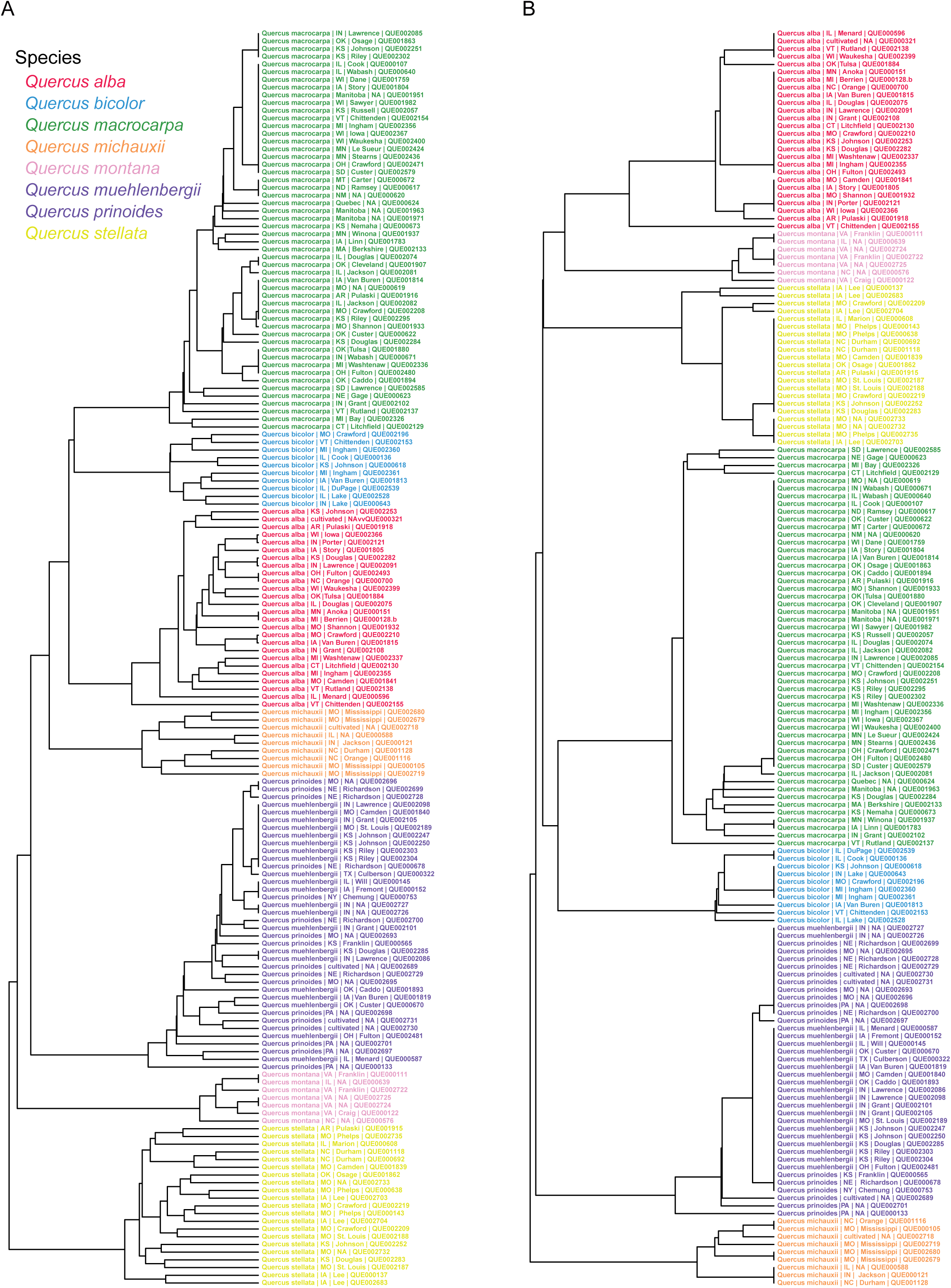
UPGMA, all loci (a) and 20 loci (b). UPGMA was conducted on a Euclidean distance matrix calculated from a three-state nucleotide matrix, where each nucleotide present for each SNP is coded as 0 = absent, 1 = 1 copy (i.e., individual is heterozygous for that SNP), 2 = 2 copies (i.e., individual is homozygous for that SNP). (A) UPGMA clustering based on all 75 loci. (B) UPGMA clustering based on 20 loci hand-selected for their decisiveness in the species sample represented here (cf. Fig. 2, red bars).

**Figure 4.**
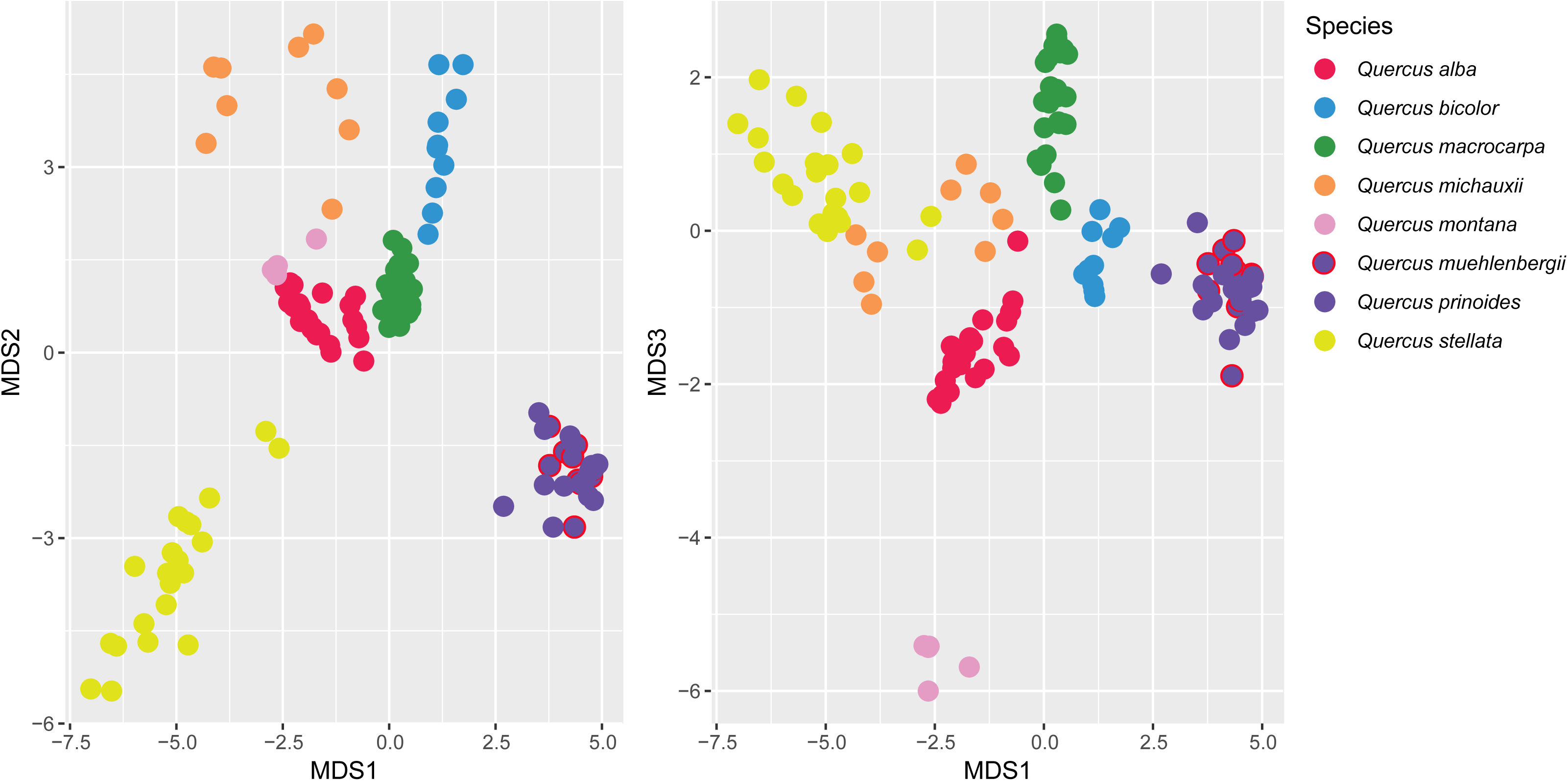
NMDS ordination, 75 loci. Non-metric multidimensional scaling was conducted in three dimensions for the same Euclidean distance matrix utilized in the UPGMA figure reported above. NMDS ordination final stress was 0.08607 and failed to reach convergent solutions in 20 iterations, but all replicate ordination attempts distinguished all pairs of species in at least one dimension, as seen in this figure.

Bayesian admixture analysis in STRUCTURE favors a *K* = 4 solution using the Δ*K* statistic of Evanno et al. (2005). Given the susceptibility of STRUCTURE and particularly the Δ*K* statistic to the highest hierarchical level of genetic structure in a dataset, we find the *K* = 4 solution not a useful description of genetic structure in our phylogenetically structured dataset. To the contrary, the *K* = 4 clustering does the best job at separating species by clade, following well supported phylogenetic relationships (Hipp *et al.*, 2018), viz. four clusters comprising *Quercus macrocarpa* and *Q. bicolor*; *Q. alba*, *Q. michauxii*, and *Q. montana*; and *Q. stellata* and *Q. muehlenbergii* / *prinoides* each on their own (Fig. 5). Given our phylogenetically structured sample, it is not surprising that Δ*K* favors a configuration that splits individuals among clades above the species level. STRUCTURE continues to distinguish species up until *K* = 8, with 7 species pairs yielding individuals admixed 10% or more based on our markers (Figs. 5, 6). Notably, it is not until *K* = 8 that the 7 species are distinguished from each other, perhaps due to high genetic variation within species that is not adequately resolved with these markers. One individual identified as *Q. alba* in the field shows evidence of introgression from both *Q. macrocarpa* and *Q. bicolor.* In the *K* = 8 configuration, *Q. bicolor* gives the appearance of being uniformly admixed with *Q. montana* at a relatively low level (9/10 individuals < 10% admixed). However, this appears to be artefactual, as the phenomenon is absent in the *K* =6, 7, and 9 configurations, all of which show genetic separation between *Q. bicolor* and *Q. montana*. In the *K* = 8 configuration, *Q. alba* resolves as a mix of two genotypes, which we combine in estimating the number of individuals admixed at 5, 10, 15, or 20% (Supplemental Table S2; Fig. 6).

**Figure 5.**
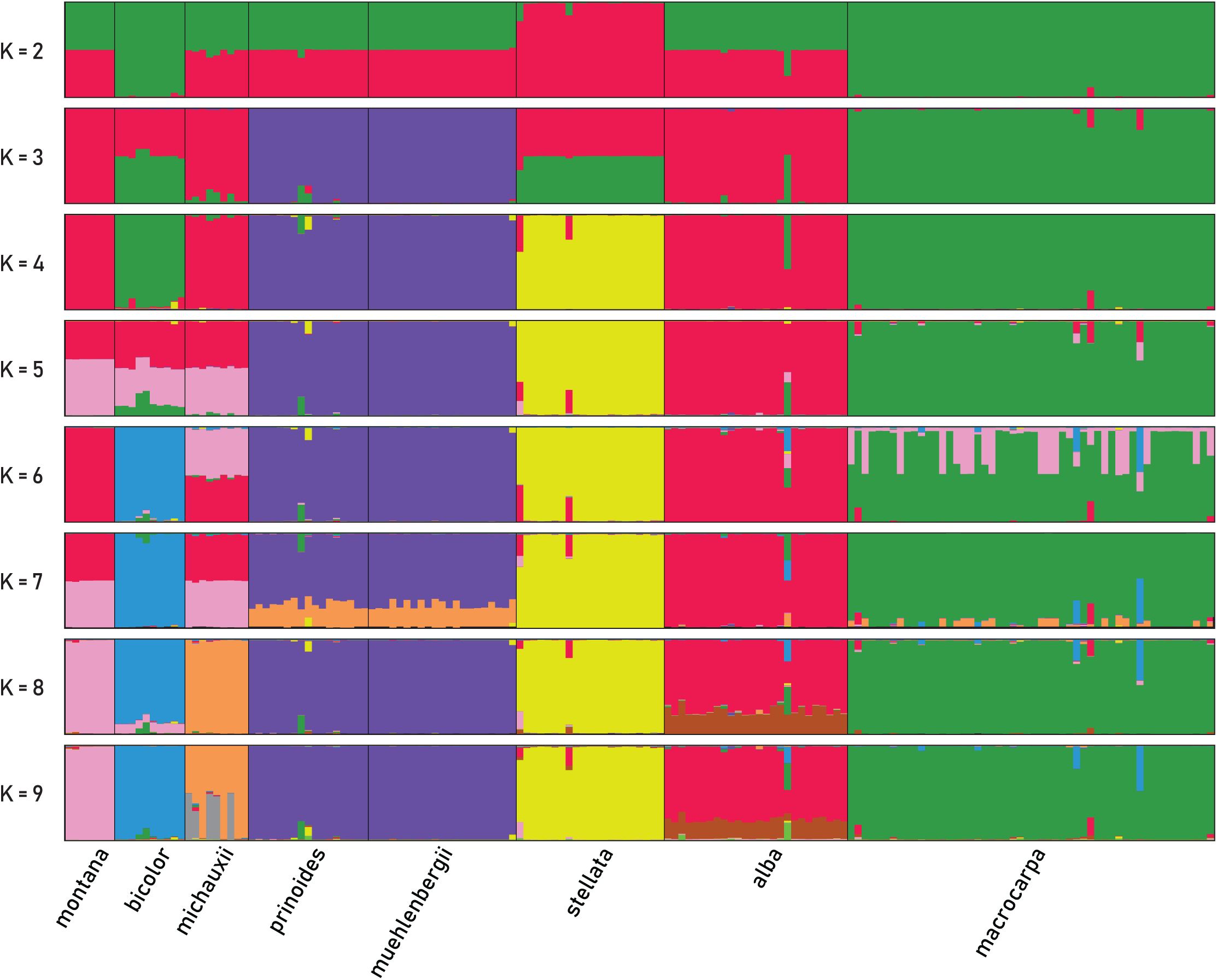
Bayesian admixture analysis conducted in STRUCTURE, assuming *K* = 2 to *K* = 9 populations. STRUCTURE analyses were conducted under the admixture model with correlated allele frequencies, from *K* = 1 to *K* =12. Values of *K* above 9 provide no additional information on population structure and are consequently not shown here. All figures represent averages over 10 independent runs of 1E06 generations each following 1E05 burn-in generations; runs were aggregated for display using the “greedy” algorithm in CLUMPP.

**Figure 6.**
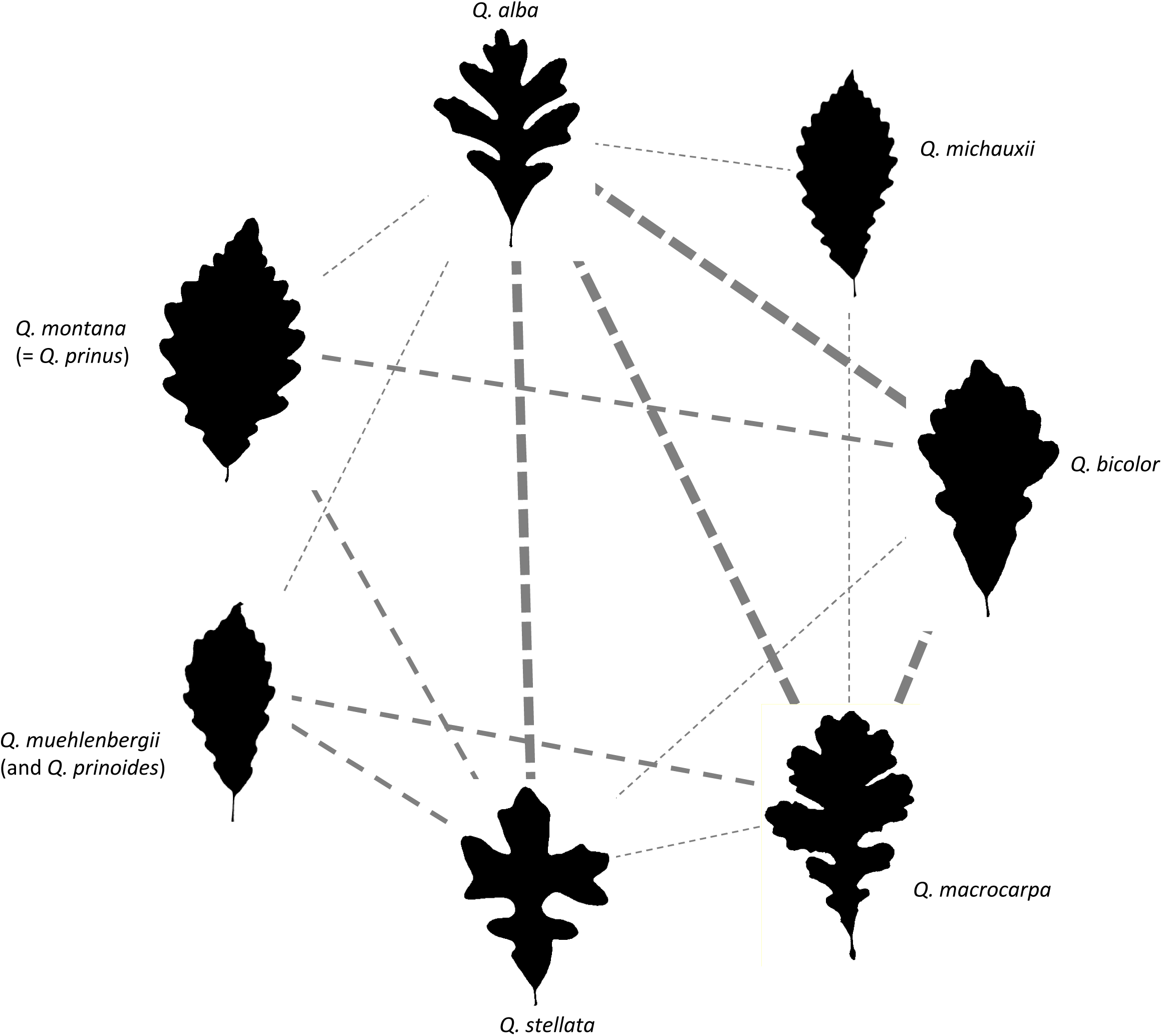
The white oak syngameon of Eastern North America *sensu* Hardin 1975, including only the species investigated in the current study. The figure replicates the 16-species figure of Hardin 1975 (his Fig. 1), including only the subset of seven species we investigated in the current study (treating *Q. muehlenbergii* and *Q. prinoides* as one), with lines indicating hybridizations that Hardin inferred from morphological study. Thin dashed lines indicate hybridizations identified by Hardin but not by us; medium dashed lines were identified by both Hardin and us, at an admixture level of 0.10 to 0.19 for at least one specimen; and thick dashed lines lines indicate admixture levels of 0.20 or higher for at least one specimen. Vouchers for leaf silhouettes are *Q. alba*: PS Manos 1838 [MOR 177669]; *Q. michauxii*: PS Manos 1843 [MOR 177659]; *Q. bicolor*: PS Manos 1847 [MOR 177662]; *Q. macrocarpa*: IL-MOR-MH108 [MOR 174544]; *Q. stellata*: PS Manos 1835 [MOR 177663]; *Q. muehlenbergii*: PM-98; *Q. montana*: PS Manos 1860 [MOR 177731].

Of the 79 RAD-seq loci used to design our SNP toolkit—79 rather than 80 because two of our SNPs derive from a single locus—59 map to a unique position on one chromosome (hereafter referred to as “uniquely mapped loci”), nine map to multiple locations in the genome, and eleven do not map to any location in the genome (Table 3; Supplemental Table S3). The uniquely mapped loci demonstrate that decisiveness is spread across the genome: 25 loci diagnostic for one or two species are found on nine out of the twelve *Quercus* chromosomes (Table 3). Moreover, distances between loci within chromosomes are mostly > 1 million bp (37 / 47 interlocus distances), and only 11% (5/47 interlocus distances) are < 10,000 bp. Distances between uniquely mapped loci averages 7.47 million bp (± 8.74 million bp, s.d.). These are all significantly clustered relative to a random draw of SNPs, under which only 0.909 interlocus distances < 10,000 bp are expected (*p* < 0.0001), 4.70 interlocus distances < 1,000,000 bp (*p* = 0.0123), and mean interlocus distance is expected to be 9.440 × 10^6^ (*p* < 0.0001). Only two of the eleven RAD-seq loci that did not map to the genome exhibit moderate decisiveness (0.81–0.869, where 1.0 or 2.0 indicates loci that are perfectly decisive for one or two species respectively). Three of the nine loci that map to multiple locations are highly decisive (1.000–1.021).

## Discussion

Our study demonstrates that with a relatively small amount of curated data—just 20 SNPs chosen to maximize genetic distinctiveness—we are able to distinguish seven genetically cohesive taxa. The fact that we are able to identify fixed or nearly-fixed SNPs across wide geographic ranges in several species suggests that introgression is distributed heterogeneously along the genome, with some areas of the genome strongly protected against introgression on a species-pair by species-pair basis. Given that these apparently-fixed SNPs are limited to our species with smallest sample size—one in *Q. alba* (N = 10), two in *Q. michauxii* (N = 9), four in *Q. montana* (N = 7)—the question of whether they are truly fixed bears further investigation. However, *Q. muehlenbergii / prinoides* is represented by 38 individuals in our dataset and three fixed SNPs, suggesting that the high-frequency proportional representation of SNPs in some species may not be an artifact of low sample size. We interpret this finding as evidence that these seven species are genetically cohesive across their ranges at least at a small number of regions of the genome, even in the face of introgression.

It is somewhat remarkable that we are able to distinguish seven interbreeding oak species with just 20 hand-picked markers. By comparison, the now-classic study demonstrating genetic distinctiveness of *Q. petraea* (Matt.) Liebl. and *Q. robur* L. utilized 20 microsatellites for just those two species (Muir *et al.*, 2000). Other studies using five (Craft & Ashley, 2006), six (Moran *et al.*, 2012), or even fifteen variable microsatellites (Aldrich *et al.*, 2003) have by contrast failed to find consistent genetic differentiation between two to three co-occuring white or red oaks (for a counter-example of relatively clean differentiation based on only 11 microsatellites, see Cavender-Bares & Pahlich, 2009). All used markers selected for variability rather than for segregation by species. Larger numbers of loci (as low as 27–28 in, e.g., Owusu *et al.*, 2015; Sullivan *et al.*, 2016) tend to pick up divergent neutral markers or markers under divergent selection (Lind-Riehl, Sullivan, & Gailing, 2014b; Sullivan *et al.*, 2016). This suggests that a moderate-sized but random sample of loci will often reflect regions of the genome that are either not yet differentiated between species (Muir & Schlötterer, 2005, 2006) or subject to ancient or contemporary gene flow (Lexer *et al.*, 2006). Because the loci that bear the stamp of population divergence history for one species pair may record introgression history for other species pairs (Crowl *et al.*, 2019; Hipp *et al.*, 2019), we would not expect any particular small set of loci to adequately describe species description across the oak phylogeny. In the current study, however, we have demonstrated that a small number *can* suffice to distinguish numerous species in a multispecies syngameon.

The SNPs we have utilized may be linked to loci under strong selection. They may as a consequence not be representative of the genome as a whole. As discussed in the paper in which these SNPs were published (Fitzek *et al.*, 2018), we selected SNPs by querying a RAD-seq dataset for loci that had pairwise F_ST_ > 0.95. Such outlier loci can tell much more refined stories about population divergence than loci that are not under such strong selection (Scotti-Saintagne *et al.*, 2004; Guichoux *et al.*, 2013; Lind-Riehl *et al.*, 2014b) and may thus pick up on divergence histories that are not clear from a broader sample of loci. These selected genes may occur in islands of differentiation distributed across the genome (Scotti-Saintagne *et al.*, 2004; Goicoechea *et al.*, 2015) and have the potential to explain genetic cohesion across species ranges even when populations diverge at neutral loci (Morjan & Rieseberg, 2004) or to differentiate species that are exchanging genes more frequently across the remainder of the genome (Lind-Riehl, Sullivan, & Gailing, 2014a; Gailing & Curtu, 2014; Oney-Birol *et al.*, 2018; Hipp, 2018). This gives them practical utility as a species identification toolkit. A genome-scale investigation, as has been conducted in the European white oaks (Leroy *et al.*, 2017, 2018), would be required to characterize the genomic architecture of differentiation among these species and address the question of whether species differences are concentrated in divergent loci under strong selection. For the time being, our study suggests that a relatively small number of selected genes may suffice to *diagnose*—not define—species, even in the face of ongoing introgression.

Despite the low sampling of loci in our study, we do find significant clustering on the genome of loci within 1 Mbp of each other (*p* = 0.008) or within 10 Kbp of each other (*p* < 0.001). This supports earlier studies that have found significant clustering of high-F_ST_ loci (Scotti-Saintagne *et al.*, 2004) as well as linkage disequilbrium (LD) among loci separated by as much as 20 centimorgans (cM) (Goicoechea *et al.*, 2015). While the *Pst*I RAD-seq loci used to design these SNPs are widespread on the genome, they are not randomly distributed, sited at higher-than-expected frequency within coding genes (Hipp *et al.*, 2019). However, our simulated distribution accounts for this, as it is drawn from the larger RAD-seq dataset from which our SNPs were developed. Thus the clustering of our SNPs appears to reflect genomic clustering of outlier loci that distinguish species of the eastern North American white oak syngameon. The causes, consequences, and scale of these genomic islands of differentiation among eastern North American white oaks bear investigation using higher sampling of individuals and loci.

We expect our power to detect complex patterns of introgression in a multispecies hybrid zone to be compromised by the low locus-sampling of this SNP toolkit (only 20 selected SNPs). Nonetheless, our study demonstrates that even without attempting to find hybrids, potentially biasing ourselves against detecting introgression, and even without employing the large numbers of loci generally favored for hybridization studies, we are able to identify introgressants involving several pairs of species from a sampling of natural populations (Figs. 5, 6). The fact that we have selected loci to be fixed or nearly fixed within species may aid in detecting first generation hybrids. At the same time, by selecting genes with high pairwise F_ST_, we effectively designed our SNPs within outlier loci (by definition, loci with higher-than-exected F_ST_), which may overestimate divergence between species and underestimate the proportion of the genome that is subject to introgression. The pairs that we found to be admixed at the 10% level for at least one individual were also found by Hardin to hybridize (Fig. 6; cf. Fig 1. in Hardin 1975). It remains to be seen using genomic markers that are not subject to the ascertainment bias in our study what the actual frequency and average percent of admixture is for these species.

## Conclusions

Oaks have been a bugbear of systematics since Darwin’s time, raising significant questions about what species are and how we can make sense of speciation in the face of ongoing gene flow (Arnold, 2016). Our work builds on studies that, in aggregate, suggest that oak species are genetically coherent across their ranges (Muir *et al.*, 2000; Hipp & Weber, 2008; Cavender-Bares & Pahlich, 2009; Hauser *et al.*, 2017) despite a history of introgression (Eaton *et al.*, 2015; McVay *et al.*, 2017a; Kim *et al.*, 2018). We concur with Hardin (1975), who wrote, “Neither Baranski (1975) nor I agree with Minckler (1965), who thinks that hybridization may mask evidence of races within white oak.”

Our study does not, however, speak to the *frequency* of hybridization, because our markers are selected for fixation or near-fixation within species. This bias may afford the markers increased utility to identify early-generation hybrids, but make them poor estimators of genome-wide rates of genetic exchange. It is important to note, in fact, that we could have told the story of introgression with a different hand-picked set of 20 or 80 SNPs: the “right” regions of the genome—by which we mean those regions that favor one particular gene-flow / genetic coherence process over another—will tell one story or the other. Both stories are embedded in the genome, and both are equally real. We cannot consequently assess Muller’s (1952) claim that “the bulk of claims of hybridity [in *Quercus*] are based upon trivial variations of the sort one may encounter in a relatively pure population of a single species.” What we can say is that the eastern North American white oak syngameon is composed of entities that most taxonomists would consider “good species.”

It is equally important to note that while our study demonstrates that there exist loci that distinguish species in the white oak syngameon across their ranges, it leaves open the question of *which* regions of the genome are responsible for species cohesion in oaks. As increasing evidence suggests that forest tree syngameons may be common, especially in the tropics (Caron *et al.*; Cannon & Lerdau, 2015; Kenzo *et al.*, 2019), the forces shapping how and the degree to which different regions of the genome capture different aspects of population divergence and gene flow history will be a central question—perhaps the central question—of tree biodiversity for the coming decade.

## Supporting information

Tables 1-3, XLSX format

## Footnotes

1 The history of Gray’s reports of hybrids is instructive. The first edition (Gray & Sullivant, 1848) included two hybrids in the genus *Quercus*, both reported to be “founded on” a single tree or individual. In the 1857 through 1862 editions (Gray, 1857, 1859, 1862), this number increased to three, which Gray described as “the following remarkable forms, by some regarded as species.” Gray’s language changes between 1848 and 1862—years flanking the publication of *Origin of Species*—from suggesting that these hybrids are mere sports to suggesting that they might be species of hybrid origin. Gray was a great supporter of Darwin and had an avid correspondence with him even before publication of *Origin* (Browne, 2010), and Gray’s change in language undoubtedly reflects a change in his view of the evolutionary implications of hybridization.

## Notes

#### Summary of Updates

Figure colors updated for clarity; SNPs mapped to genome, genomic autocorrelation tested using resampling of RAD-seq loci.

https://github.com/andrew-hipp/white-oak-syngameon

