## Supplementary figures and images for "Conserved DNA polymorphisms distinguish species in the eastern North American white oak syngameon: Insights from an 80-SNP oak DNA genotyping toolkit"

### Supplemental Figure 1

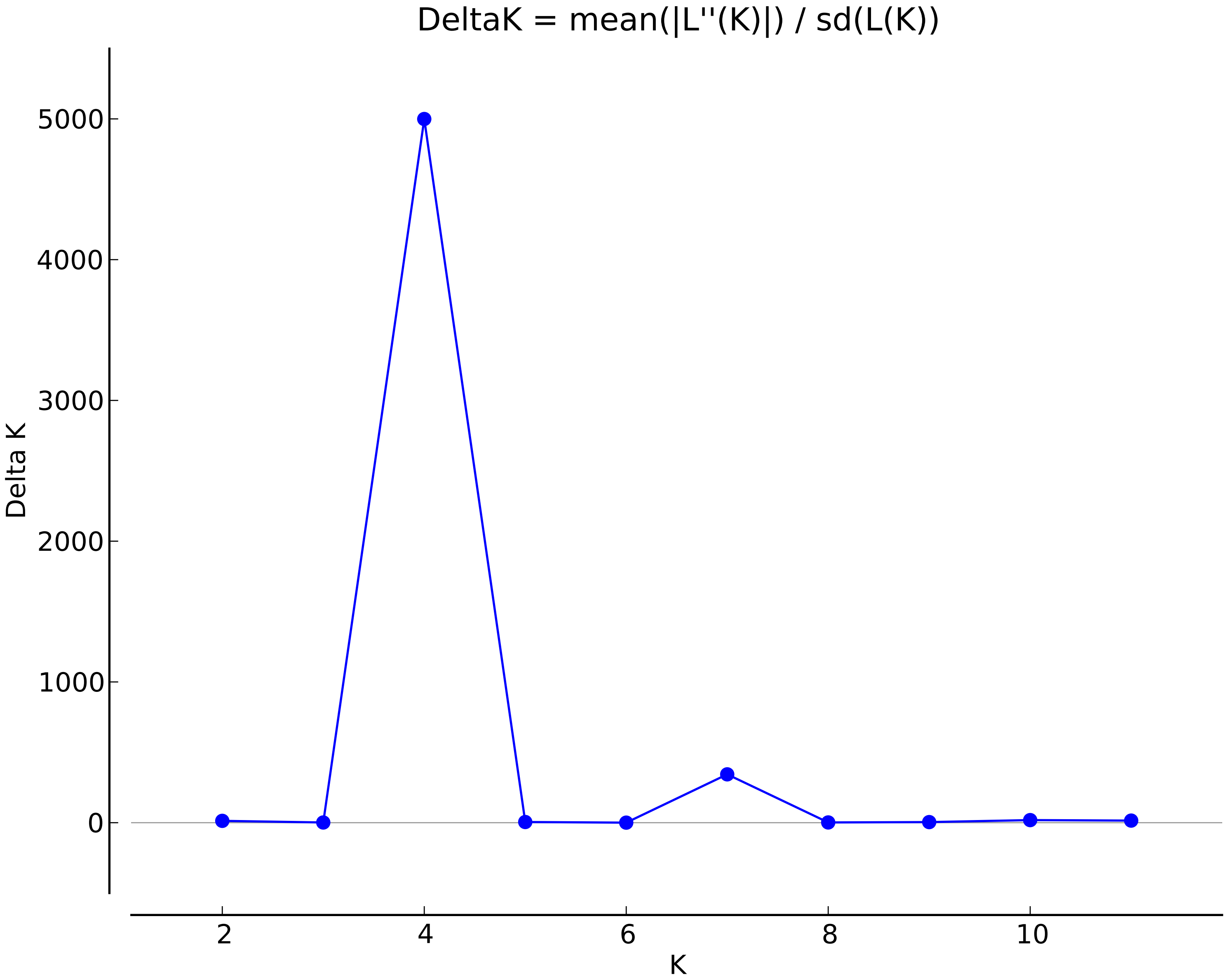
